# Temporal Metrics of Multisensory Processing Change in the Elderly

**DOI:** 10.1101/565507

**Authors:** Aysha Basharat, Jeannette R. Mahoney, Michael-Barnett-Cowan

## Abstract

Older adults exhibit greater multisensory response time (RT) facilitation by violating the race model more than younger adults; this is commonly interpreted as an enhancement in perception. Older adults typically exhibit wider temporal binding windows (TBWs) and points of subjective simultaneity (PSS) that are farther from true simultaneity as compared to younger adults when simultaneity judgment (SJ) and temporal order judgment (TOJ) tasks are utilized; this is commonly interpreted as an impairment in perception. Here we explore the relation between the three tasks in younger and older adults in order to better understand the underlying mechanisms that subserve audiovisual multisensory temporal processing. Our results confirm previous reports showing that audiovisual RT, TBWs and PSSs change with age, and we show for the first time a significant positive relation between the magnitude of race model violation in younger adults as a function of the PSS obtained from the audiovisual TOJ task with (r: 0.49, *p*: 0.007), that is absent among the elderly (r: 0.13, *p*: 0.58). Furthermore, we find no evidence for the relation between race model violation as a function of the PSS obtained from the audiovisual SJ task in both younger (r: −0.01, *p*: 0.94) and older adults (r: 0.1, *p*: 0.66). Our results confirm previous reports that i) audiovisual temporal processing changes with age; ii) there is evidence for distinct neural networks involved in simultaneity and temporal order perception; and iii) common processing between race model violation and temporal order judgment is impaired in the elderly.

## Introduction

The central nervous system (CNS) is constantly presented with information from multiple modalities that must be efficiently combined in order to form a coherent representation of the world. There is an evolutionary advantage to integrating sensory information from multiple modalities as it allows the observer to respond to external events more quickly and accurately relative to solely processing unisensory information (Stein & Stanford, 2008). One important factor that the CNS must consider when determining whether to bind multisensory information is the relative timing of events. Studies have shown that there is a window in time within which multisensory events are judged to have occurred simultaneously. Interestingly, a growing body of research has shown that this temporal binding window (TBW) changes throughout early development (Lewkowicz, 1996; Hillock et al., 2011; Hillock-Dunn and Wallace; 2012), injury (Wise & Barnett-Cowan, 2018), disease (Chan et al., 2015) and aging (Poliakoff et al., 2006; Setti et al., 2014; Bedard & Barnett-Cowan, 2016; Basharat et al., 2018). With respect to aging, a variety of experimental approaches have been utilized to characterize changes in the perceived timing of multisensory events for old compared to young adults. Here we seek to assess the relationship among some of these approaches in order to better understand the underlying mechanisms that may subserve them.

A classic psychophysical method used to assess the relative perceived timing of multisensory events is response time (RT), in which the observer is presented with unisensory or multisensory stimuli and asked to press a response key as fast as possible following stimulus presentation. Early work conducted by Raab (1962) suggested that the presentation of a pair of stimuli initiated a detection race wherein the winner’s time determined the observed RT. The race model inequality (RMI) proposed by Miller (1982) tests whether the observed RT facilitation for multimodal stimuli is too large to be attributed to statistical facilitation; RMI has become the standard testing tool in many multisensory studies. A violation of this inequality indicates that separate processing of the stimuli is not taking place and indicates synergistic neural mechanisms. Research comparing MSI effects in young and older adults is limited, however, it appears that older adults demonstrate greater multisensory RT facilitation effects compared to younger adults when presented with multimodal stimuli (Laurienti et al., 2006; Peiffer et al., 2007; Diedrich et al., 2008; Mahoney et al., 2011;Couth et al., 2017). The test of the RMI tends to show significant violations in older adults suggesting integration of the unisensory stimuli while younger adults tend to show reduced or no violation suggesting minimal integration (Laurienti et al., 2006; Mahoney et al., 2011). More recently, however, it has been argued that determining multisensory integration based on only RT differences is likely insufficient and that multisensory and unisensory cumulative distributed functions (CDFs) should be examined (Couth et al., 2017; Mahoney et al., 2018a, Mahoney et al., 2018b). As this method of assessment is not common, research comparing the amount of area under the curve (AUC) obtained from the CDF difference wave from younger and older adults can provide further information regarding age-related alterations of multisensory integration.

In the literature, multiple tasks can be found that are commonly utilized to assess multisensory processing. During the sound-induced flash illusion (SiFi), participants are presented with two auditory beeps alongside a visual flash and they are asked to report the number of perceived flashes. The illusion is induced when two visual flashes are perceived (instead of one) when two beeps and a single flash are presented in close temporal proximity (Shams et al., 2005). Using this illusion, Setti and colleagues (2014) found that older adults were maximally susceptible to the SiFi at a stimulus onset asynchrony (SOA) of 70 ms and were no longer susceptible at the SOA of 270 ms. These results indicate that this particular illusion is likely an underestimation of multisensory integration effects as it is an insensitive measure of the window of time during which individuals integrate cues from multiple modalities. In the stream/bounce illusion, a two-dimensional visual display is used to present two identical objects moving toward one another, coinciding, and moving apart (Sekuler et al., 1997). After the point of coincidence, the movement of the objects can be interpreted as if they continued in their original direction or as if they bounced off one another and reversed directions. A brief beep is presented 150 ms before or after or at the point of coincidence which increases bounce perception compared to the control condition in which no beep is presented. Previously, Roudaia and colleagues (2013) showed that older adults did not have an increased perception of the bounce when the auditory stimulus was presented at the point of coincidence - suggesting an age-related reduction in multisensory integration. Bedard and Barnett-Cowan (2016) on the other hand found that older adults were susceptible to the illusion indicating that they were integrating auditory and visual cues over a large window of time. Like the SiFi, one of the concerns with the stream/bounce illusion is that it is not sensitive to the full parameterization of the temporal window during which multisensory information is integrated and it only provides an indirect method of assessing such a window (Sekuler et al., 1997). Simultaneity judgment (SJ) and temporal order judgment (TOJ) are extensively used in the literature (Allan, 1975; Mitrani et al., 1986; Vatakis et al., 2008; Barnett-Cowan & Harris, 2009; Barnett-Cowan & Harris, 2011; Love et al., 2013) as they are sensitive to both TBW, a window of time during which information from multiple sensory modalities is bound and perceived as synchronous, as well as the point of subjective simultaneity (PSS), the point at which participants are most likely to indicate simultaneity. In these tasks, participants are presented with pairs of multisensory stimuli and are asked to determine whether the two stimuli were presented simultaneously or which stimulus came first, respectively. Although they both provide measures of the TBW and PSS, previous research has shown that these two tasks measure different perceptual processes (Mitrani et al., 1986; Vatakis et al., 2008; Barnett-Cowan & Harris, 2009; Barnett-Cowan & Harris, 2011; Love et al., 2013) and are likely to be subserved by different neural mechanisms (Allan, 1975; Dhamala et al., 2007; Adhikari et al., 2013; Linares & Holcombe, 2014; Basharat et al., 2018). Using these tasks, it has been found that the TBW varies with age as it tends to be wider in early childhood, becomes more fine-tuned during middle childhood, and widens again with aging (Lewkowicz, 1996; Poliakoff et al., 2006; Hillock et al., 2011; Hillock-Dunn and Wallace; 2012; Setti et al., 2014; Bedard & Barnett-Cowan, 2016). A wider TBW in older adults indicates that they are more likely to perceive synchrony and thus have more trouble differentiating temporally offset stimuli (Virsu et al., 2003; Poliakoff et al., 2006; Hay-McCutheon et al., 2009; Alm and Behne, 2013; Busey et al., 2010; Chan et al., 2014a; Chan et al., 2014b). Widening of the TBW with aging is of concern given that information that should be encoded as arising from separate events is more likely to be integrated, which can result in decreased speech comprehension (Maguinness et al., 2011; Setti et al., 2013), an inability to dissociate from distracting or inaccurate information (Wu et al., 2012), and an increased susceptibility to falls (Setti et al., 2011; Mahoney et al., 2014; but see Mahoney et al., 2018a) and fall awareness (Lupo & Barnett-Cowan, 2017). Furthermore, age-related impairments in driving performance and speech comprehension have been associated with temporal processing deficits within the auditory (Gordon-Salant & Fitzgibbon, 1993; Babkoff & Fostick, 2017) and visual (Wood, 2002; Lacherez et al., 2014) domain. In order to address this concern, psychophysical training regimes have been designed to recalibrate the TBW which may prevent these deficits from developing in the older population (Powers et al., 2009; Chan et al., 2014a, Chan et al., 2014b).

SJ, TOJ, and RT are different methods of assessing temporal perception of events, however, no study to date has compared the three tasks. This comparison is paramount as it provides us with a better understanding of how multisensory information is processed and whether or not there is a relation between the different decision-making processes that underlie the behaviour associated with these tasks. In this study, we aim to explore the relation between the three tasks in younger and older adults in order to better understand the underlying mechanisms that subserve multisensory temporal processing and change with age. Previously research on the relation between RT and TOJ was assessed and revealed that the TBWs obtained from the RT task were wider than for the corresponding TOJ task (Diederich & Colonius, 2015; Mégevand et al., 2013); this can be seen as an observer’s strategy to optimize performance as the TOJ task requires participants to discern small asynchronies in which a narrower window is beneficial, whereas a wider window would maximize multisensory facilitation (as determined by the race model violation) in the RT task. Diederich and Colonius (2015) suggested that these ideas can be further expanded to the SJ task, whereby a relation between the TBW and RT should be observed. Furthermore, if a relation between the width of the TBW and RT exists, one could argue for such a relation between RT and PSS as well, primarily as longer RTs would provide the CNS with a better opportunity to determine the PSS more accurately. To assess such a relation in the aging population, some researchers have varied the stimulus onset asynchronies (SOAs) used in the RT task to extract the TBW. Here, older adults have slower RTs compared to younger adults, they also have a broader TBWs, and they tend to show greater multisensory facilitation as assessed via race model violation (Diederich et al., 2008; Laurienti et al., 2006).

These studies provide initial evidence for a relation between a wider TBW, slower RT, and a larger violation of the race model. However, research comparing RMI (quantified as the AUC from the CDF difference wave) to measures obtained from SJ and TOJ (i.e., TBW and PSS) in younger and older adults is lacking and can reveal further information regarding the underlying mechanisms that are involved in decision making. As multisensory processing changes with age (i.e., requiring light to appear much earlier than sound to perceive simultaneity as compared to younger adults), assessing these relations within this population provides further information related to whether the underlying mechanisms associated with these tasks maintain their relation.

The main objective of the current study is to determine age-related differences in auditory-visual (AV) integration using a unique experimental design that encompasses aspects of AV RT, SJ, and TOJ. Here, we aimed to determine age-related differences in TBW, PSS, and the magnitude of race model violation. We hypothesize that 1) older adults will have slower RTs as compared to younger adults; 2) older adults will demonstrate larger race model violations compared to younger adults; 3) increased race model violations will be positively correlated with wider TBWs; and that 4) increased race model violations will be positively correlated with PSS falling farther away from true simultaneity.

## Materials & methods

### Participants

Participants (n=56) were recruited from the University of Waterloo (younger adults) and from the Waterloo Research in Aging Participant Pool (WRAP; older adults). The WRAP program ensures that all recruited participants are healthy older adults over the age of 60 with no significant medical concerns (i.e., Alzheimer’s disease, Parkinson’s disease, stroke, epilepsy, etc.) by screening for mild cognitive impairment and dementia using the Montreal Cognitive Assessment (MoCA; mean score = 27, SD = 0.47. A score ≥ 26/30 indicates normal cognition; Nasreddine et al., 2005) and reviewing self-reported data acquired from medical history questionnaires.

Male and female participants between 19 and 79 years of age were included in this study. Participants included 30 young (17 females, mean age = 22.93, s.e. = 0.66) and 26 older adults (19 females, mean age = 70.80, s.e. = 0.90). All participants were required to have normal or corrected-to-normal vision and hearing. Prior to study inclusion, participants completed a selfreported clinical information form where they indicated (yes/no) if they had normal or corrected-to-normal vision and if they had normal or corrected-to-normal hearing. If participants answered no to any of the above questions, they were subsequently excluded from the study. In appreciation of their participation, participants received a $10 per hour remuneration. This study was approved by the University of Waterloo’s Human Research Ethics Committee in accordance with the Declaration of Helsinki, and written informed consent was obtained from all participants before participation.

### Experimental setup

Each participant completed three experimental tasks while seated in front of a 23.6 inch ViewSonic V3D245 computer monitor (resolution 1920 × 1080, 120 Hz) in a sound-proof booth with his/her head stabilized on a chin rest. Visual stimuli were presented on the monitor at a viewing distance of 57cm, in the form of white circles (0.4°). Auditory stimuli were emitted from two speakers (Altec Lansing Multimedia computer speaker system, ACS95W) adjacent to the monitor such that they were 66 cm apart. A Macbook Pro (OS 10.9 Mavericks) that resided outside of the booth was used to run the tasks. VPixx Technologies ProPixx hardware and DataPixx software version 3.01 was utilized for this experimental procedure to ensure the synchrony of the audio and visual stimuli (depending on condition) with <1 ms accuracy. Participants were able to record their response for each trial by using the RESPONSEPixx handheld 5-button response box.

### Procedure

Participants completed the SJ, TOJ, and RT tasks in a randomized order. For all tasks, a central fixation cross (visual angle = 0.5°) was presented on the screen, and participants were instructed to fixate this cross throughout the experimental procedure. In order to reduce temporal predictability, each trial began with the stimulus being presented after a delay of 1000 - 3000 ms. Participants were presented practice trials prior to commencement of each of the experimental tasks. Test performance during the actual experiment was monitored on a laptop from outside the booth.

### Simultaneity judgment

In the SJ task, participants were instructed to report, using different response buttons, whether they perceived the auditory and visual stimuli as occurring simultaneously (right button) or not (left button). Participants were explicitly told to respond as accurately as possible as opposed to responding quickly. Visual stimuli were presented in the form of a 0.4° white circle (49.3 cd/m^2^) against a black background (0.3 cd/m^2^), which appeared 2° below the fixation cross for 17 ms. They were either preceded or followed by an auditory beep (1850 Hz, 7 ms, 71.7 dB) at the following SOAs: 0, 25, 50, 100, 150, 200, 300 ms. Ten trials were presented in a randomized order for each condition for a total of 130 trials (see Figure 1).

**Figure 1.**
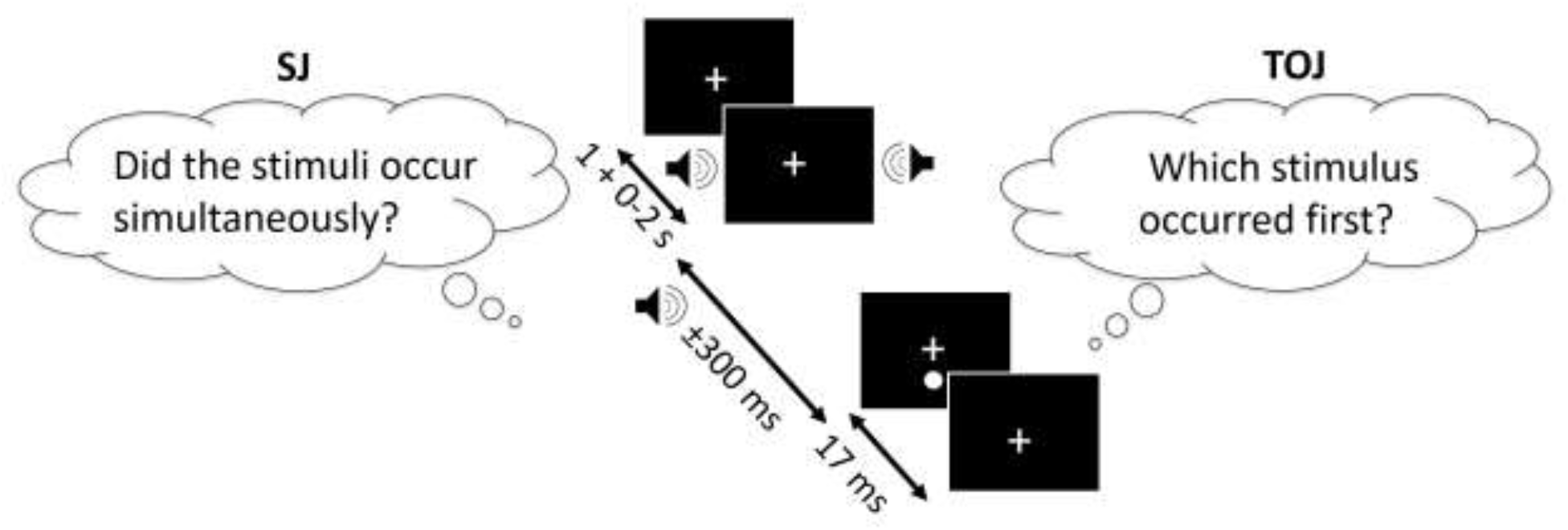
SJ task (left) and the TOJ task (right), presented with the SOAs of 0, ±25, ±50, ±100, ±150, ±200, ±300 ms (-ve = sound appeared before light). In both tasks, the first stimulus of the audiovisual pair can appear 1-3 sec following the fixation cross and the second stimulus appears between 0 – 300 ms after the first stimulus. The figure depicts the auditory stimulus (i.e., beep) as presented before the visual stimulus (i.e., flash). Note, that the experimental design for the SJ and TOJ is identical, however the instructions vary by task.

### Temporal order judgment

The experimental design of the TOJ task was identical to the SJ task with the exception of the task instructions. Here, participants were asked to report, using the response buttons, whether they perceived the visual (right button) or the auditory (left button) stimulus as appearing first; “synchronous” or “I don’t know” responses were not acceptable for this task. Again, participants were explicitly told to respond as accurately as possible as opposed to responding quickly.

### Reaction time task

In the RT task, participants were told that they would either see a flash of light, hear a beep, or a combination of the two. Participants were instructed to press the response button as soon as they detected any one of the three experimental conditions: unisensory Visual (V), unisensory Auditory (A) or multisensory (AV). In order to maintain consistency across all three tasks, the same stimuli and durations were employed. Each stimulus was presented 100 times in a random order (300 trials in total). Trials were divided into 2 blocks and participants were given a break in between blocks to reduce fatigue (see Figure 2).

**Figure 2.**
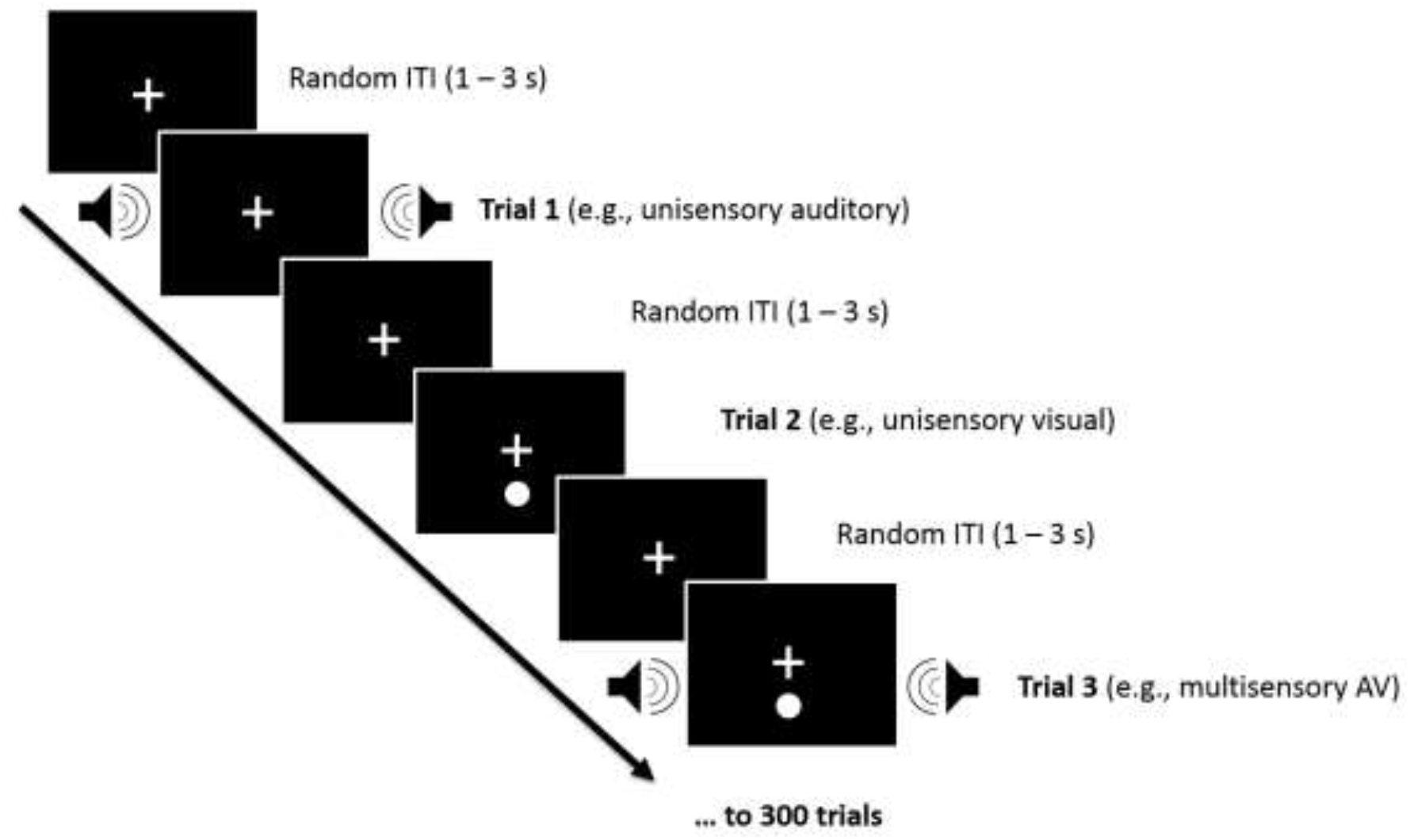
Participants were presented with unimodal [auditory (A) or visual (V)] or bimodal [audiovisual (AV)] stimuli and were asked to make speeded responses to all stimuli, regardless of sensory modality by pressing a RESPONSEPixx button which triggered the next trial. Two blocks of A, V, and AV stimuli (150 trials per block) were randomly presented with random inter-trial-intervals (ITIs) of 1 – 3 s.

## Statistical analysis

### Simultaneity and temporal order judgment tasks

To estimate the accuracy (PSS values) and the precision (TBW) with which participants made their judgments for SJ and TOJ, psychometric functions were fitted to participant’s responses as a function of SOA using SigmaPlot version 12.5. Each task was analyzed individually for each participant, with participant data fit to both Gaussian (for the SJ task; Eq. 1) and logistic (for the TOJ task; Eq. 2) functions:

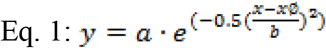

Where *a* is the amplitude, *xø* is the PSS and *b* is the standard deviation.

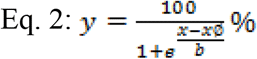

Where *a* is fixed to 1, *xø* is the PSS and *b* is the standard deviation (slope between 0.25 and 0.75).

The best fit parameters corresponding with the PSS and TBW were identified for each participant separately, and those participants whose data was poorly estimated were excluded from further analysis (r^2^ ≤ 0.2; YA = 1; OA = 3).

As we are interested in the relationships between TBWs obtained from the two tasks and not their absolute size, we chose to analyze the *b* values (i.e., standard deviation) of these psychometric functions as a proxy for the size of the TBW to avoid discrepancies in the literature that differ when defining the absolute size of the TBW.

Using a within-subjects design, paired t-tests were conducted to assess differences between TBWs and PSSs within each group. Independent t-tests were used in accordance with Leven’s test for equality of variance to further assess differences between younger and older adults. Pearson’s correlations (α = 0.05) were assessed between the two tasks for all participants while controlling for age. Furthermore, Pearson’s correlations (α = 0.05) were conducted to determine age-specific relations.

### Reaction time task

#### Error analysis, outlier removal, and mean RT analysis

As previously mentioned, participants responded to 300 trials in total (100 per condition). Given recent reports implicating the use of RMI over RT facilitation for investigating MSI effects (Couth 2017 & Mahoney et al 2018a, Mahoney et al 2018b), we applied a similar approach. RMI was first tested using Gondan’s permutation test over the fastest quartile (0-25%) of responses and violation was indeed observed for both younger (t_max_ = 4.42, t_crit_ = 2.21, *p* < 0.001) and older adults (t_max_ = 5.71, t_crit_ = 2.08,*p* < 0.001) (Gondan, 2010; Gondan & Minakata, 2016). Data trimming procedures were not applied (see Gondan, 2010, 2016; Mahoney, 2018a; Mahoney, 2018b); however slow responses and misses (defined as > 1500 ms or not registered by the program respectively [< 3% for each condition]) were set to infinity rather than excluded (see also Mahoney & Verghese, 2019 for a RMI tutorial). To be consistent with other MSI studies, RT facilitation (multisensory condition – most efficient unisensory condition) was also calculated.

#### Mean RT analysis

A 2 (age group: young or old) x 3 (condition: auditory, visual, or audiovisual) repeated-measures RM ANOVA was conducted to determine whether age and condition significantly affected RT. Mauchly’s test of sphericity was conducted and the Greenhouse-Geisser was used if necessary. Planned pairwise comparisons were also made to assess the differences between young and older adults by condition.

#### Test of the race model

As previously mentioned, the race model posits that the response to redundant signals is produced by the modality that processes its respective signal the fastest and thus is the “winner” of the race (Raab, 1962). Race model violations are typically tested using cumulative distribution function (CDF) models which compare the actual CDF distribution to the predicted CDF distribution (Miller, 1982).

Each participant’s data was sorted in ascending order for all three conditions (A, V, AV). Each participant’s RTs were then quantized into 5^th^ percentile bins until the 100^th^ percentile was reached, yielding 21 bins in total.

*Actual* CDF distributions were formed using the following equation (Eq. 1):

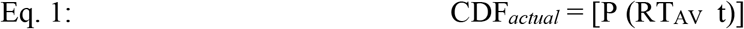

Where RT_AV_ represents the RT observed for the multisensory condition for any latency, t (Colonius & Diederich, 2006; Mahoney et al., 2011).

*Predicted* CDF models were formed using the following equation (Eq. 2):

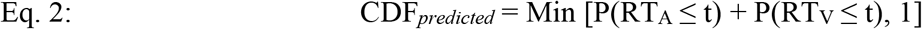

Where RT_A_ and RT_V_ represent the RTs observed for unisensory condition A (e.g., auditory) and V (e.g., vision), for any time, t (Colonius & Diederich, 2006; Mahoney et al., 2011).

Differences between the *actual* CDF distribution and the *predicted* CDF distribution were calculated for every participant across all percentile bins as follows (Eq. 3):

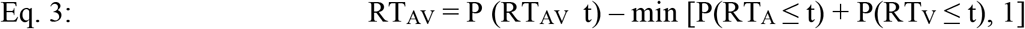

When the *actual* CDF is less than or equal to the *predicted* CDF the race model is accepted. However, the race model is violated when the *actual* CDF is greater than the *predicted* CDF. Thus, a negative value (or zero) indicates acceptance of the race model while values greater than zero provide evidence for multisensory integration as they are indicative of race model violation (Colonius & Diederich, 2006; Mahoney et al., 2014).

Although many researchers have previously utilized t-tests (i.e., paired t-tests comparing actual vs. predicted CDF or one sample t-tests along the difference curve) to determine race model violations, it has been argued, that these tests are too conservative (Gondan & Minakata, 2016). As mentioned above, we used a data-driven approach to determine RMI violations by conducting Gondan’s permutation test over the fastest quartile (0-25%) of responses, where robust violations were evident for both younger and older adults (see also Figure 7 below). In order to compare the AUC obtained from younger and older adults, a Kruskal-Wallis test was conducted in order to account for the non-normal distribution of the AUC data.

### Relation between the TBW and the race model

Furthermore, in order to assess the relation between race model violations, SJ, and TOJ, Pearson’s correlations (α = 0.05) were determined between the AUC values, TBWs, and PSSs for younger adults. While Spearman’s correlations (α = 0.05) were determined between the AUC values, TBWs, and PSSs within older adults.

## Results

### Simultaneity and temporal order judgment tasks

Participants’ responses were fitted to either a Gaussian or a sigmoidal logistic curve for SJ and TOJ respectively using equation 1 or 2 from which PSS and TBWs were extracted for analysis. Figure 3 shows the average Gaussian functions (SJ) and Figure 4 shows the average logistics function (TOJ) for younger and older adults. The goodness of fit from the SJ task for younger (r^2^ Mean: 0.85, Median: 0.88, SD: 0.09, s.e.: 0.02) and older adults (r^2^ Mean: 0.80, Median: 0.84, SD: 0.13, s.e.: 0.03) were similar [independent t-test: *t*(55) = 1.68, *p* = 0.1)]. The goodness of fit from the TOJ task for younger (r^2^ Mean: 0.76, Median: 0.82, SD: 0.18, s.e.: 0.03) and older adults (r^2^ M: 0.75, Median: 0.81, SD: 0.21, s.e.: 0.042) were also similar [independent t-test: *t*(53) = .53, *p* = 0.6)].

**Figure 3.**
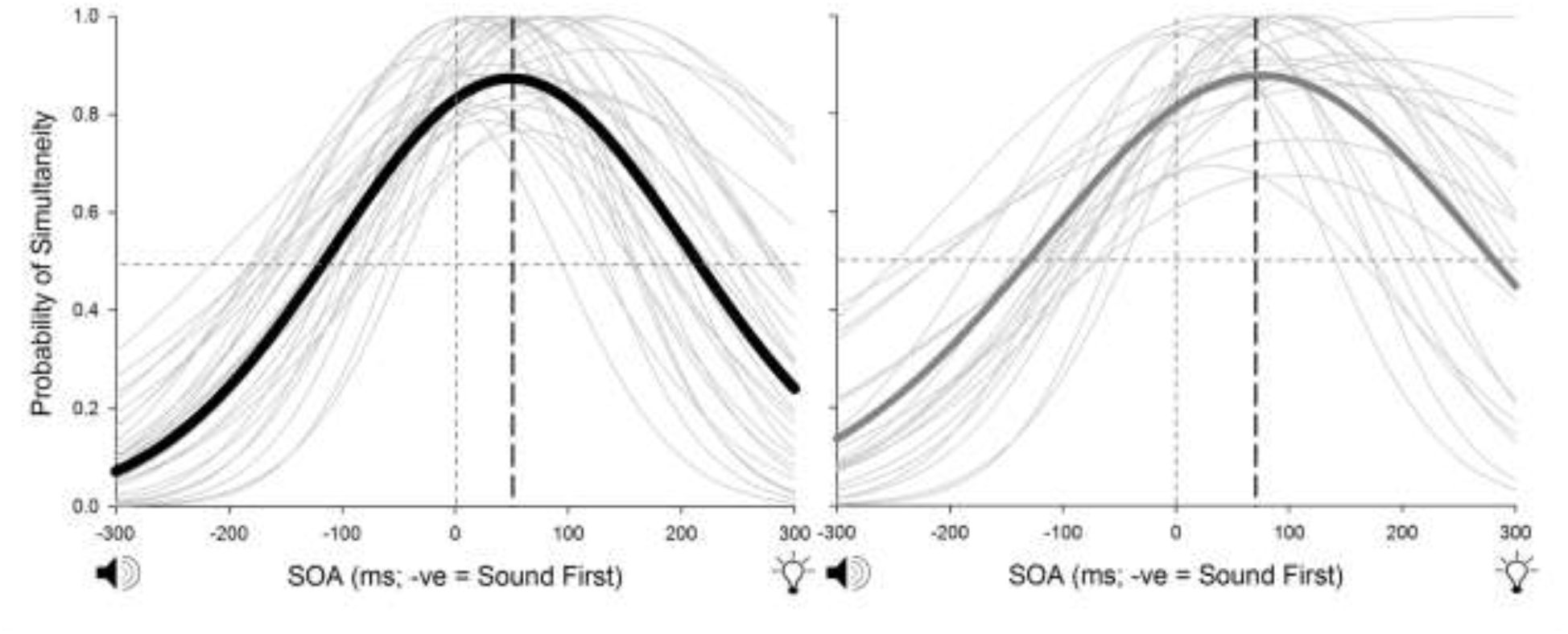
SJ: here the Gaussian function is fit to the average (thick lines) and individual (thin lines) data. Younger adults (black) require the visual stimulus to occur approximately 58 ms before sound while older adults (grey) require the visual stimulus to occur approximately 82 ms before sound in order to perceive the two stimuli as simultaneous.

**Figure 4.**
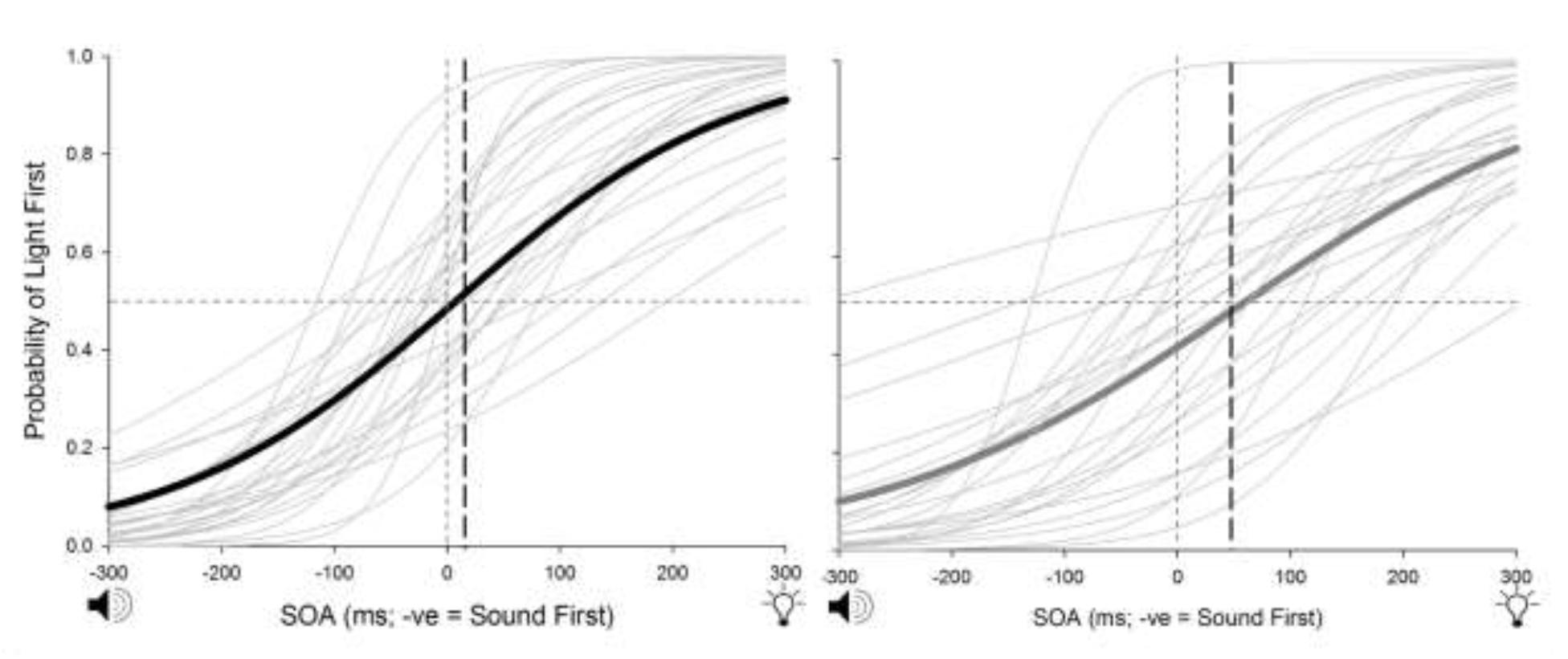
TOJ: the sigmoidal function is fit to average (thick lines) and individual (thin lines) data. Here, younger adults (black) require the visual stimulus to appear approximately 10 ms before sound while older adults (grey) require the visual stimulus to appear approximately 49 ms before light in order to perceive the two stimuli as being simultaneous.

In line with our previous work (Basharat et al., 2018), within the younger group, the paired t-test revealed that the TBW obtained from the SJ task (M = 160.07, s.e. = 8.39) was significantly wider than the TOJ task (M = 103.04, s.e. = 10.51); (*t*(27) = 6.69,*p* < 0.001; Figures 3 and 4). Furthermore, a paired t-test between the two tasks for the PSS revealed that the visual stimulus was required to appear before the auditory stimulus earlier in the SJ task (M = 56.36, s.e. = 7.77) than the TOJ task (M = 10.31, s.e. = 14.17) in order for simultaneity to be perceived; (*t*(28) = 2.81, *p* < 0.01; Figures 3 and 4). Within the older group, like the younger adults, a paired t-test revealed that the TBW was wider in the SJ task (M = 186.26, s.e. = 14.35) compared to the TOJ task (M = 117.01, s.e. = 13.12); (*t*(22) = 4.74,*p* < 0.001; fig 3 and 4). No significant difference was found between the PSS for the SJ task (M = 82.70, s.e. = 9.36) compared to the TOJ task (M = 53.91, s.e. = 23.52) within the older group; (*t*(22) = 1.16, *p* = 0.26; Figures 3 and 4).

Planned independent t-tests were conducted on each task to determine age-related differences between the two tasks. While the test did not reach significance, older adults (M_SJ_ = 186.26, s.e. = 14.35; M_TOJ_ = 117.01, s.e. = 13.12) exhibited wider TBWs compared to younger adults (M_SJ_ = 162.59, s.e. = 8.48; M_TOJ_ = 103.04, s.e. = 10.51) for both SJ (*t*(50) = -.95, *p* = 0.35) and TOJ (*t*(49) = -.84, *p* = 0.40). As predicted, no significant effects of PSS were found between the younger (M_SJ_ = 56.35, s.e. = 7.76; M_TOJ_ = 10.31, s.e. = 9.25) and older adults (M_SJ_ = 79.26, s.e. = 14.17; M_TOJ_ = 53.91, s.e. = 23.51) for SJ (*t*(50) = −1.84, *p* = 0.07) and TOJ (*t*(50) = −1.66, *p* = 0.10).

Age controlled partial correlations were first conducted on all participants for the TBW as well as the PSS. In line with previous literature (Bedard & Barnett-Cowan, 2016) a significant positive correlation was found between the TBWs obtained from both the tasks (*r*(49) = 0.50, *p* < 001). No significant correlations were observed for PSS (*r*(49) = 0.03, *p* = 0.84). Pearson’s correlations were then conducted within each group and the TBWs from the two tasks within both younger (*r*(28) = 0.61, *p* < 0.001) and older (*r*(23) = 0.44, *p* = 0.04) adults were found to be significantly positively correlated. PSSs were not correlated between the two tasks in both younger (*r*(29) = −0.04, *p* = 0.85) and older adults (*r*(23) = 0.06, *p* = 0.77).

### Reaction time task

#### Error analysis, outlier removal, and mean RT analysis

Both young and older adults made few errors with an overall accuracy of 99.98% and 99.99% in each group respectively. Younger adults maintained an accuracy of 99.4% in the auditory trials, 99.6% in visual stimuli, and 97.9% in the audiovisual trials. Older adults achieved an accuracy of 99.6% in auditory trials, 99.8% in visual trials, and 98.0% in audiovisual trials. In line with Couth and colleagues (2017) our data also revealed most outliers to be the slower responses (>1500 ms) with very few misses (<1% for all conditions). Outliers were converted to infinity and only correct responses were included in the analyses.

#### Mean RT analysis

Results from the 2 (age group: younger, older) x 3 (condition: audio, visual, audiovisual) RM ANOVA revealed a significant main effect of group (*F*(1, 25) = 25.97,*p* < 0.001; η_p_^2^ = 0.51; Figure 5) and condition (*F*(1.30, 32.4) = 129.37,*p* < 0.001; η_p_^2^ = 0.84). The interaction between group and condition was also significant (*F*(1.54, 38.48) = 21.49,*p* < 0.001; η_p_^2^ = 0.46). In line with our hypothesis, planned pairwise comparisons revealed that older adults (M = 369.26 ms, s.e. = 17.68) demonstrated significantly longer RTs compared to younger adults (M = 280.85 ms, s.e. = 14.03; *p* = 0.001). The pairwise comparisons also revealed that responses to audiovisual trials (276.35 ms, s.e. = 12.02) were significantly faster than auditory (324.80 ms, s.e. = 17.45) and visual trials (374.00 ms, s.e. = 11.23;*p* < 0.001; see Figure 5).

**Figure 5.**
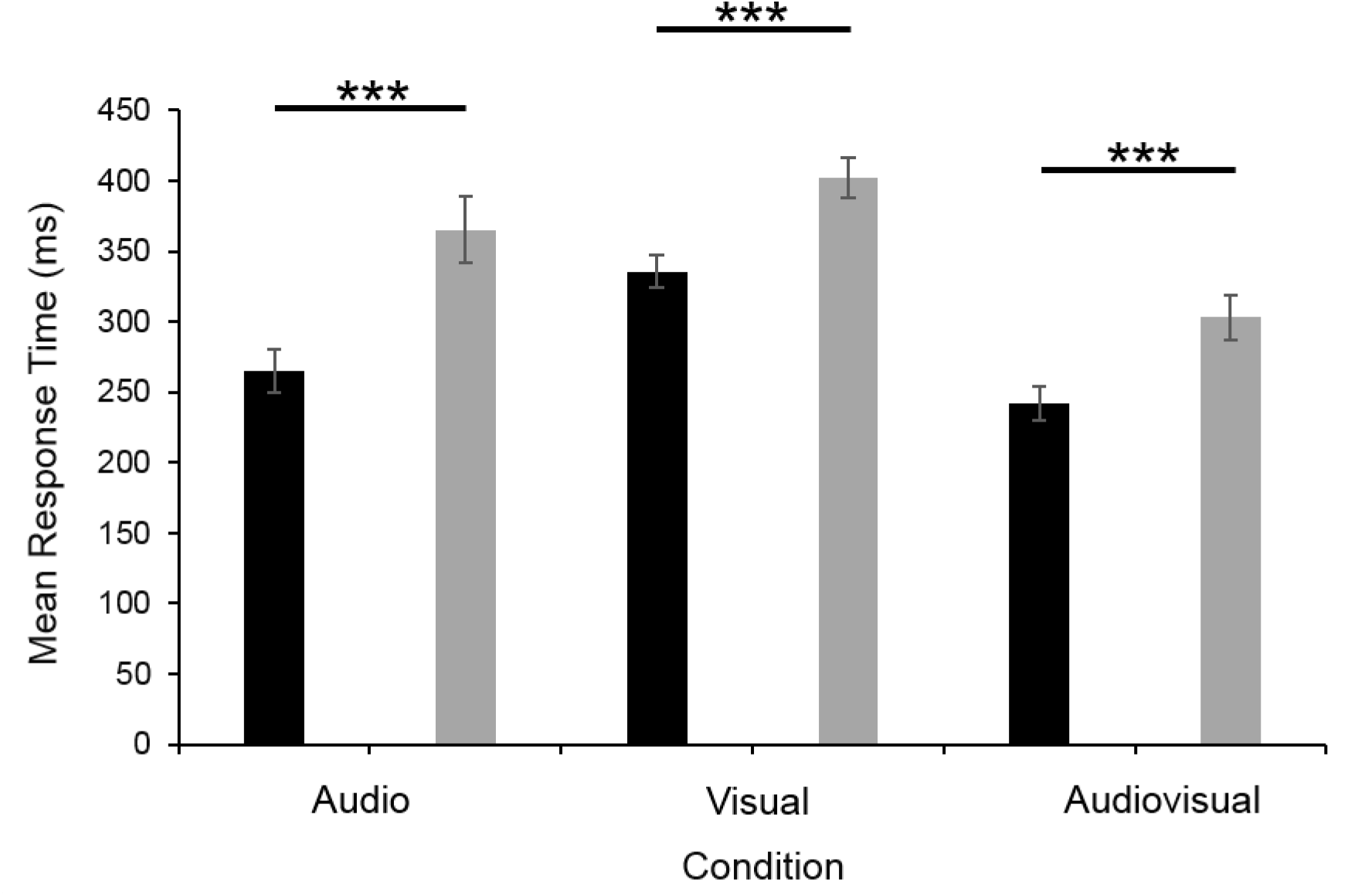
Averaged response time data (with SEMs) from the auditory, visual, and audiovisual conditions for both younger (black) and older (grey) adults. *Error bars* are ± 1 SEM. *Asterisks* indicate statistical significance at *p* < 0.001 level.

#### The race model violation

The difference waveform, calculated by subtracting the predicted CDF from the actual CDF, is indicative of whether or not the race model has been violated (Colonius et al., 2006). Evidence for the co-activation model and thus support for multisensory integration is provided if a positive value is obtained regardless of the significance of the magnitude (Colonius et al., 2006; Mahoney et al., 2011, 2015; see Figures 6 and 7). Figure 7 indicates a violation of the race model and provides evidence for the co-activation model over the first 25^th^ percentile (highlighted in grey) in both younger and older adults. These findings are consistent with the main effect of condition (i.e., audio, visual, audiovisual) as found through the RM ANOVA conducted above with mean RT. However, as mentioned above, Gondan’s permutation test was also conducted to statistically assess race model violations and was significantly violated in both younger and older adults. We then conducted a Kruskal-Wallis test comparing the AUC values to determine group differences; here, the AUC data obtained from older adults violated the Shapiro-Wilk test due to two outliers; D(23) = 0.87, *p* < 0.01. A statistically significant difference between the groups was determined χ^2^ (1) = 8.48,*p* < 0.01 where younger adults showed a smaller mean rank score (21.05) compared to older adults (33.37) thus indicating larger race model violations in older adults (see Figures 6 and 7).

**Figure 6.**
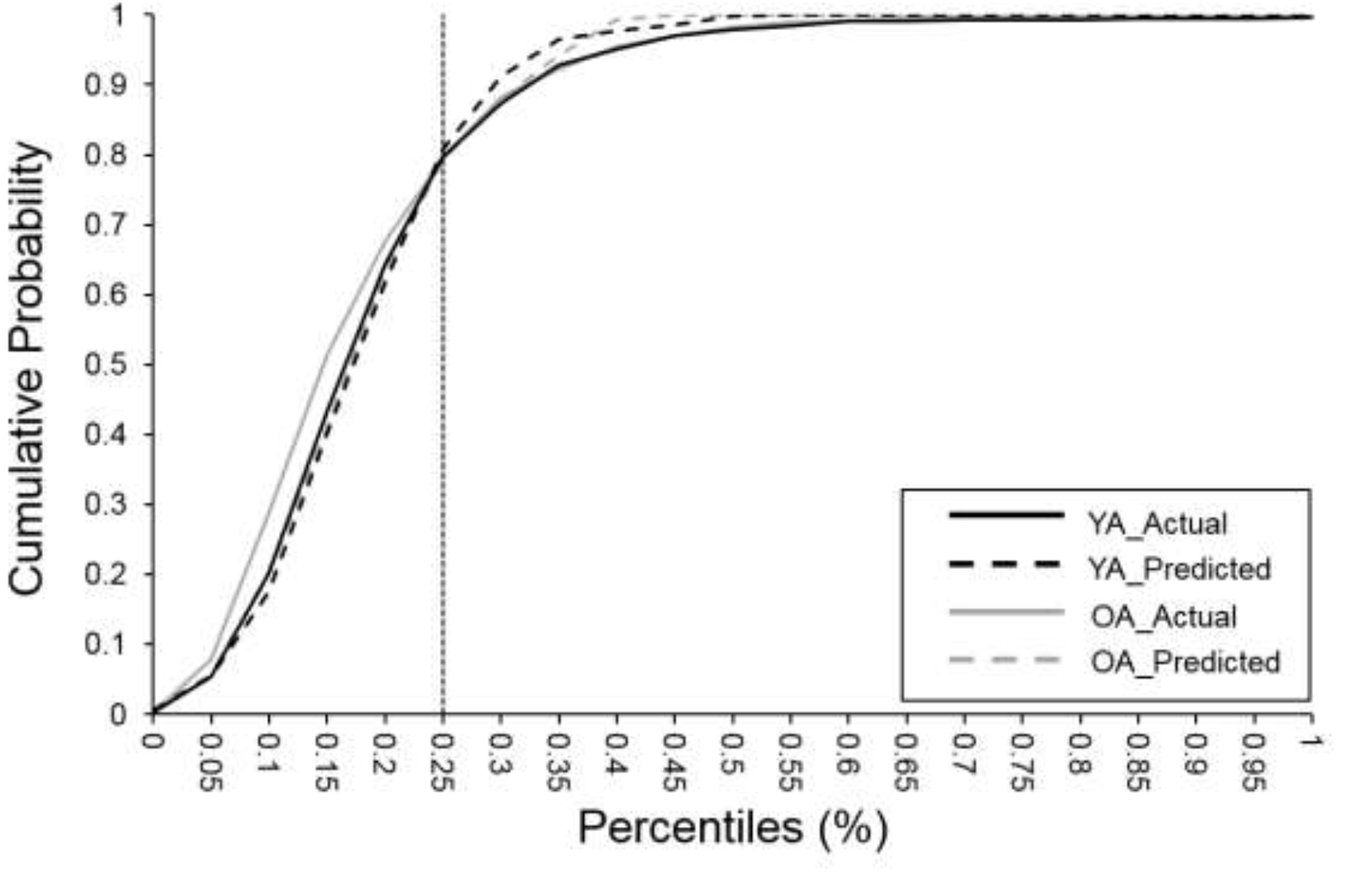
Cumulative probability graphs where the solid lines represent the actual cumulative probability while the dashed lines represent the predicted cumulative probability for younger (black) adults and older (grey) adults.

**Figure 7.**
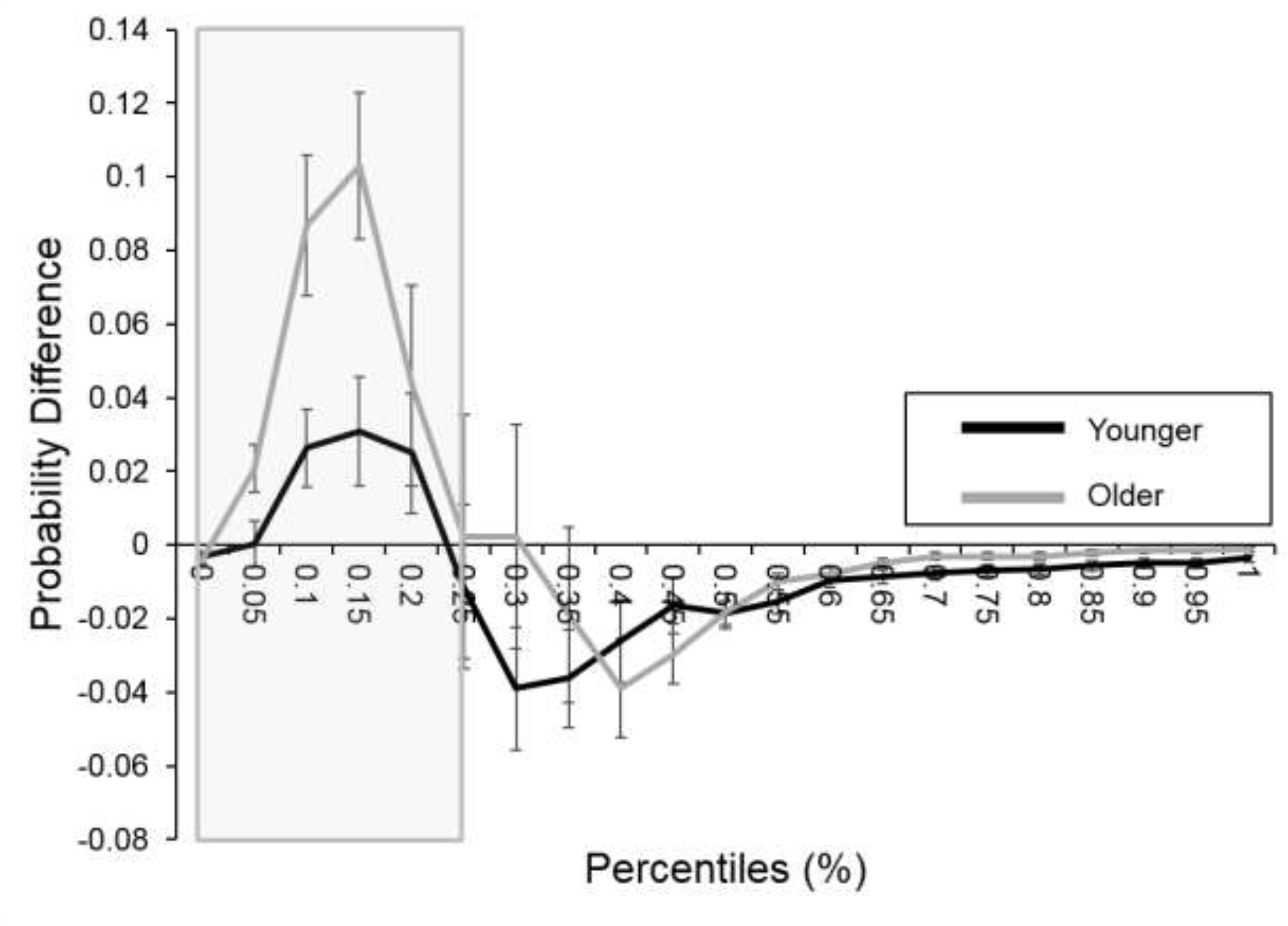
Test of the race model by group. The graph represents the probability difference wave where the predicted CDF is subtracted from the actual CDF for younger (black line) and older (grey line) adults. They grey box indicates the area over which the analyses were conducted. *Error bars* are ± 1 SEM.

### Relation between SJ, TOJ, and RT tasks

Prior to conducting the correlation analysis, data was checked for normality. All data was normally distributed except for the AUC values obtained from older adults which consisted of 2 outliers. Thus, Pearson’s correlations (α = 0.05) were determined between the PSSs and the TBWs obtained from the SJ and TOJ tasks with the average AUC values obtained from the RT task over the fastest (0-25%) percentiles within the younger adults while Spearman’s correlations (α = 0.05) were conducted for older group. A significant positive correlation was found between the PSS obtained from the TOJ task and the AUC (r(29) = 0.49,*p* < 0.01) in the younger group. No other correlations were found for both the younger and older group (see Figures 8 and 9).

**Figure 8.**
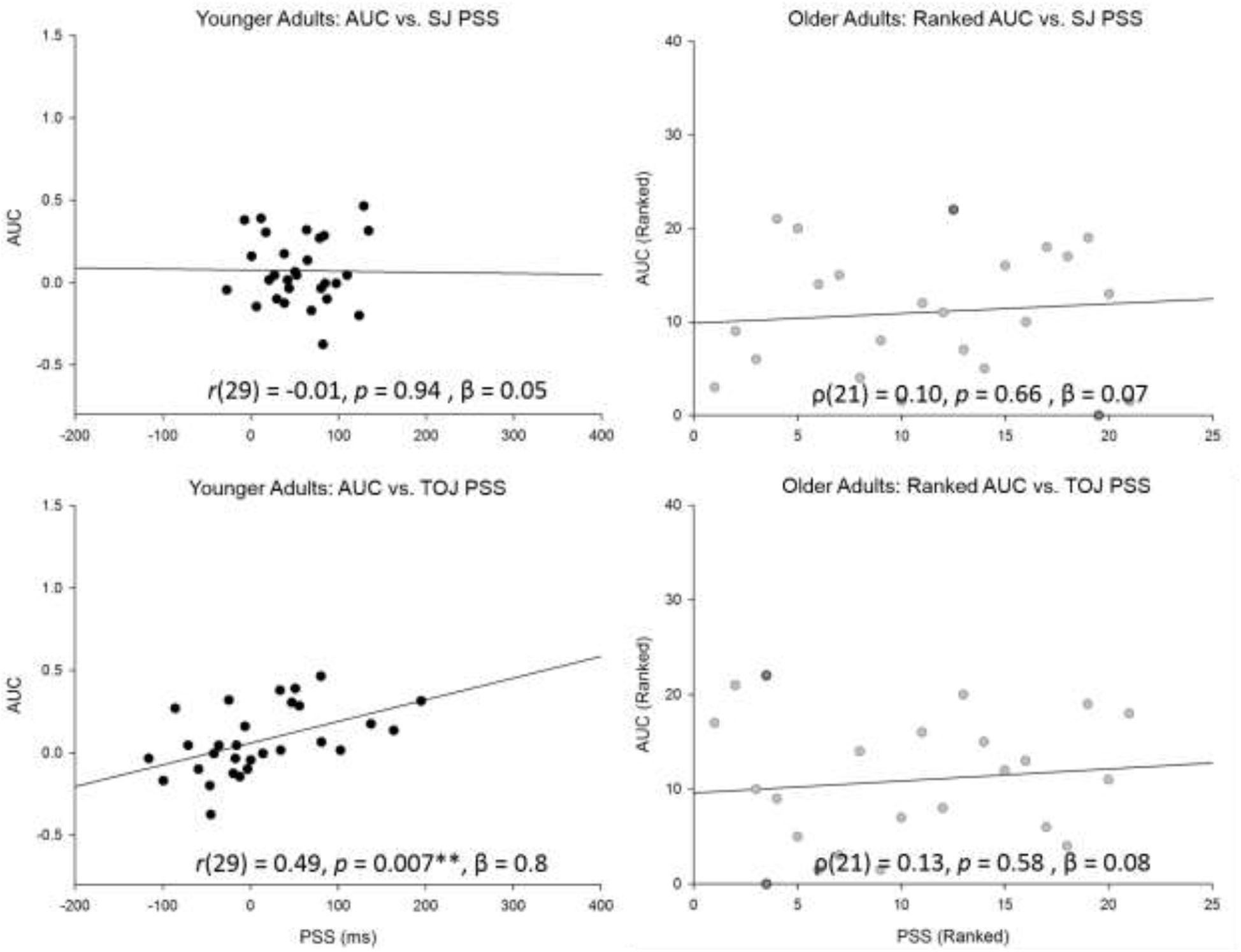
Scatter plots of correlations between the PSS and AUC for both younger (left) and older (right) adults. Notice that Spearman’s correlations were conducted for older adults and thus the ranked data is reported for the group; note the two outliers (dark grey) found within the older adult group. Pearson’s correlations were conducted for younger adults. Note: only the PSS obtained from the TOJ task is positively correlated with the AUC. ** P < 0.01.

**Figure 9.**
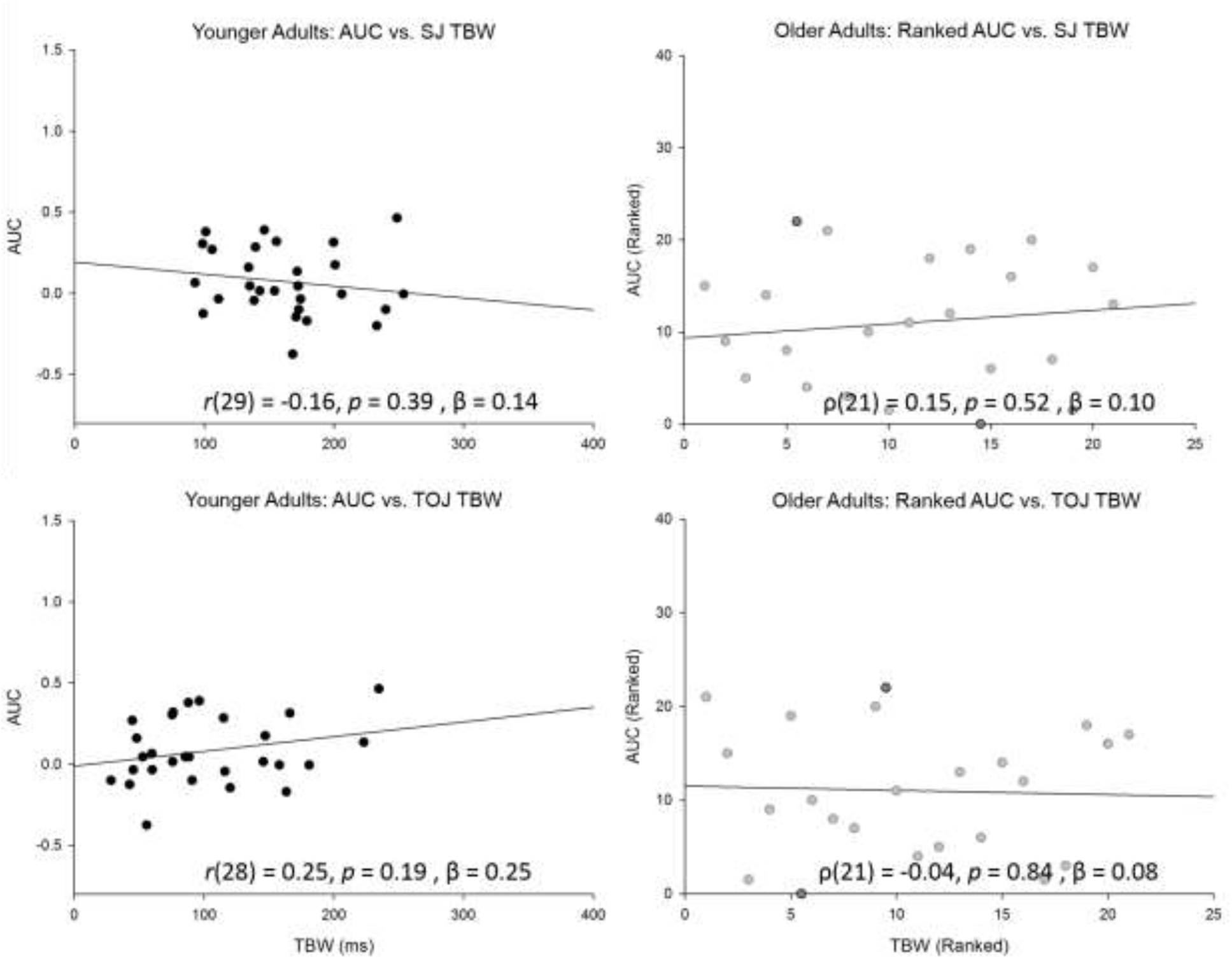
Scatter plots of correlations between the TBW and AUC for both younger (left) and older (right) adults. Notice that Spearman’s correlations were conducted for older adults and thus the ranked data is reported for the group. Notice the two outliers (dark grey) found within the older adult group. Note that Pearson’s correlations were conducted for younger adults.

## Discussion

The main objectives of this study were to identify whether relations exists between race model violation (as assessed via AUC) and measures obtained from SJ and TOJ tasks (i.e., PSS and TBW). We found a highly significant positive correlation between race model violation and the PSS from the TOJ task. Surprisingly, there is no evidence of an association between the AUC and the PSS for the SJ task in younger adults. Figure 8 clearly shows that within the younger group, those who do not violate the race model are more likely to require the sound to be presented before light in order to perceive the two as being simultaneous. Whereas those that require the light to be presented before sound in order to perceive simultaneity are more likely to violate the race model. However, for SJ, we found that all but one individual require light to be presented prior to sound in order to perceive simultaneity and yet approximately 60% violate the race model while the remaining 40% do not. This suggests that there is a discrepancy regarding the underlying mechanism such that race model violation is related to audiovisual integration and perception for TOJ but not for SJ. This is in line with previous literature which suggests that SJ and TOJ are subserved via different neural mechanisms and therefore differ from one another (Adhikari et al., 2013; Dhamala et al., 2007; Setti et al., 2011a; Basharat et al., 2018).

What do our results reveal regarding changes in multisensory processing among the elderly? The very strong positive correlation that was found between AUC and PSS from the TOJ task for younger adults disappeared within the older group. This may be related to the fact that temporal order perception seems to be impaired with aging while simultaneity perception may be preserved (Bedard & Barnett-Cowan, 2016). We speculate that if a relation between PSS and AUC was found for the SJ task in younger adults, such a relation would have persisted with aging and would also have been found within the older group. Not only do we not find such a relation, we also find two non-integrators in the older adult group who demonstrate that some older adults fail to integrate audiovisual information. This is quite interesting given that older adults tend to benefit more from multisensory integration compared to their younger counterparts (Laurienti et al., 2006; Peiffer et al., 2007; Diedrich et al., 2008; Mahoney et al., 2011; Couth et al., 2017), but perhaps is not surprising given Mahoney and colleagues recent reports of differential multisensory integration patterns for older adults (2014, 2015, 2018a, 2018b). Correlation between race model violation and PSS were determined without the outliers, however, the relation did not reach significance.

SJ, TOJ, and RT tasks are three of the most common tasks utilized in the literature to assess multisensory integration however no other study has compared measures obtained from all three tasks (i.e., TBW, PSS, and RMI). We argue that understanding the relation between race model violation, as assessed via the RT task, and TBW and PSS, as assessed via SJ and TOJ tasks, can provide further information regarding MSI and the underlying mechanisms that may change with age. Interestingly, literature from the SJ and TOJ tasks argues that there is an **impairment** in the elderly’s ability to perceive the temporal order of events from multiple modalities due to an increase in the width of the TBW (i.e., less precision) and a larger shift from true simultaneity (i.e., less accuracy; Poliakoff et al., 2006; Setti et al., 2011a, b; Chan et al., 2014 a, b; Bedard & Barnett-Cowan, 2016). Whereas studies testing RT argue that there is a greater **enhancement** in performance (i.e., faster RTs) for multimodal stimuli especially in the aging population (Laurienti et al., 2006; Peiffer et al., 2007; Diedrich et al., 2008; Mahoney et al., 2011; Couth et al., 2017). This suggests that different decision-making processes may be at play for the two categories of tasks (i.e., SJ/TOJ vs. RT).

What might explain these similarities and differences in temporal metrics of multisensory processing? We know from early research that the superior colliculus is implicated in processing both unisensory (i.e., audio or visual) and multisensory (i.e., audiovisual) stimuli for simple RT tasks (Meredith & Stein, 1983, 1986; King & Palmer, 1985). We also know from more recent research that the superior colliculus is also involved in assessing temporal order of auditory and visual cues (in addition to the superior colliculus, the posterior parietal cortex, the superior temporal sulcus, and frontal cortices) (Calvert et al., 2001; Ghazanfar & Schroeder, 2006). However, we find differences in cortical activation between simple RT and temporal order perception as well. From other literature using simple RT tasks, we also find evidence for early multisensory convergence in cortical areas that were previously considered as being ‘unisensory’ (Giard & Peronett, 1999; Molholm et al., 2002; Schroeder & Foxe, 2005). For example, using a simple RT task, Giard and Peronnet (1999) found ERP activation representing multisensory processing as early as 40 ms post-stimulus presentation over the occipital region indicating that multisensory integration takes place much earlier than expected. Forced-choice paradigms exploring synchrony and asynchrony perception on the other hand provide evidence for activation in higher order regions. For example, in a study conducted by Dhamala and colleagues (2007), participants were asked to judge whether audiovisual stimuli were presented simultaneously, whether a sound was presented first, a light was presented first, or if they could not tell. They found that the primary visual sensory cortices, parietal, and prefrontal cortices are involved in asynchrony perception. The left temporal and parietal cortices, as well as the right frontal cortex and superior colliculus are involved in synchrony perception (Adhikari et al., 2013; Dhamala et al., 2007). Our results align with previous research indicating that although there is a relation between race model violation and asynchrony perception, such a relation may not exist for synchrony perception (as indicated by a lack of relation between RT and SJ). The different neural mechanisms that underlie the behaviour observed for an RT task and synchrony perception may explain why no such relation is found with aging. It is important to note however, that some behavioural models posit that perception (as measured by SJ and TOJ) and automatic response (as measured by RT) may be subserved via similar internal mechanisms (Cardoso-Leite et al., 2007; Miller & Schwarz, 2006; Sternberg & Knoll, 1973).

It has previously been reported that a wider TBW is associated with slower RT (Laurienti et al., 2006; Poliakoff et al., 2006; Diederich et al., 2008), however we failed to find such an association. Furthermore, although not significant, older adults showed wider TBWs compared to younger adults for both the SJ and TOJ tasks. This may explain the lack of relation between TBW and the AUC in older adults as we had predicted that older adults would have much wider TBWs and would also violate the race model more so than younger adults. It is important to note that this lack of difference between younger and older adults has been seen before. Previously, with a similar sample size, Basharat and colleagues (2018) yielded similar results where older adults exhibited wider TBWs compared to younger adults but they were not statistically significant. Whereas Bedard and Barnett-Cowan (2016) found that older adults had significantly wider TBWs compared to younger adults on the TOJ task but not on the SJ task. This finding suggested that simultaneity perception may be preserved with aging while temporal order perception is not. As the study conducted by Bedard and Barnett-Cowan (2016) had a larger sample size and was well-powered, we speculate that the larger sample size contributed to the significant differences observed for the TBW values between the two groups. In line with previous literature, we did not find a relation between the two groups for the PSS values (Bedard & Barnett-Cowan, 2016; Basharat et al., 2018).

In agreement with previous literature, the mean analysis of the reaction time data indicates that older adults had significantly longer RTs compared to younger adults regardless of modality and that providing stimuli from multiple modalities significantly decreased response time (Laurienti et al., 2006; Peiffer et al., 2007; Mahoney et al., 2011). Furthermore, providing evidence for our hypothesis, the results showed that although both the younger and older group violated the race model, older adults were more likely to do so. Different theories have been put forward to explain such improvements in multisensory integration in the elderly. One possible explanation is the principle of inverse effectiveness; it states that reduced sensitivity in the individual sensory systems (i.e., decreased visual acuity (Spear, 1993), increased auditory thresholds (Liu and Yan, 2007), and decreased olfactory capabilities (Rawson, 2006)) combined with age-related alternations in cognitive processing (i.e., decline in executive function (Falkenstein et al., 2006), working memory, and attention (Fabiani, 2012)) increases the magnitude of multisensory enhancement (Hariston et al., 2013; Freiherr et al., 2013). Mozolic and colleagues (2012) have provided another explanation for the improvement observed in the older group; they state that older adults do not adequately filter sensory noise and hence are more prone to distractions compared to younger adults. However, as the background sensory information becomes more relevant, older adults benefit from enhanced processing of such information. It is clear that the neural networks involved in multisensory integration change with age and these alterations directly impact multisensory processing in the aging population. Using magnetoencephalography, Diaconescu and colleagues (2013) compared neural activity of younger and older adults to unimodal and multimodal audiovisual stimuli and found that younger adults showed increased activity in sensory-specific regions after multimodal stimuli were presented whereas older adults showed activity in the inferior parietal and medial prefrontal areas. These results provide evidence for posterior to anterior shift with aging (PASA) indicating that older adults engage frontal brain areas to a greater extent than younger adults in order to compensate for impaired function in other brain areas (Grady et al., 1994; Davis et al., 2008). Age-related changes clearly have large implications on multisensory processing and this study has provided further evidence that older adults benefit more from multimodal cues.

Given that poor multisensory processing has been correlated with speech comprehension deficits (Maguinness et al., 2011; Setti et al., 2013), an inability to dissociate from irrelevant information (Wu et al., 2012), and poor driving performance (Ramkhalawansigh et al., 2016), measures of MSI present an easy assessment tool to be utilized in the clinical setting. However, prior to the inclusion of these tasks in the clinic, it is important for researchers to understand the relation between them. Our results indicate that there is a relation between the point at which participants perceive simultaneity and the likelihood of violating the race model which may change with age. Knowing this information suggests that the ‘impairment’ and the ‘enhancements’ observed here may be subserved by similar mechanisms, but further research is required to untangle why these differences arise as we age. One explanation for the differences observed between younger and older adults may be related to a general cognitive decline due to structural changes and loss of brain mass (Mozolic et al., 2012). However, if general cognitive decline could completely explain the differences in performance between the two groups, older adults would consistently perform poorly regardless of whether unimodal or multimodal cues were presented. As indicated by our results and previous research, older adults demonstrate greater multisensory enhancement from bimodal cues compared to younger adults and thus age-related changes cannot fully be explained by general cognitive slowing (Peiffer et al., 2007). Another explanation for why such differences arise may be associated with age-related changes in gamma-aminobutyric acid (GABA), the principal inhibitory neurotransmitter in the CNS (Takayama et al., 1992; Gao et al., 2013; Porges et al., 2017). Previous research has found approximately a 5% reduction in GABA concentration per decade of aging after adolescence in the frontal cortex leading to a decline in inhibitory signals. This reduction in GABA has been associated with an inability to inhibit binding of erroneous cues, thereby resulting in an increased inability to determine temporal order of stimuli (Goa et al., 2013). In addition to between group differences, our results also indicate a large variation in multisensory perception within each group. In the future, this inter-individual variability can be further investigated through genetic factors which may contribute to the heterogeneity of the results and may explain the differences observed in younger and older adults.

## Conclusion

Here, we have demonstrated that older adults are impaired in judging temporal order and simultaneity, due to an extended TBW. However, older adults also exhibit greater enhancement in performance on the RT task as indicated by a higher likelihood of race model violation. Correlations conducted to assess the relation between the three tasks reveal that the likelihood of violating the race model is associated with the point at which simultaneity is perceived but only for the TOJ task. No such relation was found in the older group. By utilizing RT, SJ, and TOJ, our work provides further evidence that the underlying mechanisms that subserve these tasks change with age. Future studies should attempt to determine the underlying neural mechanisms that subserve these three tasks and to develop training paradigms that increase the accuracy and precision with which the elderly bind multisensory information in order to reduce errors in temporal order and simultaneity judgments.

## Acknowledgements

This work was generously supported by an Natural Sciences and Engineering Research Council of Canada (NSERC) Discovery Grant (#RGPIN-05435-2014) to MB-C.

